# Preferential innervation of endometriosis by hyperexcitable Ret/GFRα1+ nociceptors associates with target GDNF and clinical pain

**DOI:** 10.64898/2026.08.21.744503

**Authors:** Adam J. Dourson, Makenna Fluegel, Avery J. Kim, Juliet M. Mwirigi, Maria Elena P. Morales, Ruben Borja, Judith P. Golden, Elise Bardawil, Whitney T. Ross, Hadas Nahman-Averbuch, Robert W. Gereau

## Abstract

Endometriosis is a prevalent condition characterized by chronic pelvic pain that is frequently refractory to treatment. While the mechanisms underlying this pain remain poorly defined, clinical evidence often indicates that lesion innervation, but not disease stage (e.g. number and depth of lesions), correlate with pelvic pain severity. However, characterization of lesion-innervating neurons is incomplete, revealing an opportunity to identify novel, disease-modifying therapeutics. Here, we coupled functional analyses of lesion-innervating neurons in a mouse model with concurrent identification and characterization of lesion-innervating neurons from pain-phenotyped endometriosis patients. Following the confirmation of abdominal-directed pain-like behaviors in the mouse model, electrophysiological analysis revealed that lesion-innervating dorsal root ganglion (DRG) neurons are hyperexcitable compared to matched controls. These neurons are predominantly small-diameter and bind Isolectin B4, an established marker of the GDNF Family Ligand receptor, Ret. GDNF is concentrated within the stromal layer of both mouse and human lesions, adjacent to axons expressing the GDNF co-receptor, GFRα1. Critically, clinical pain correlates with lesion GDNF level, axonal density, and neuronal GFRα1 levels. These data provide evidence that endometrial lesions may recruit the Ret-positive subpopulation of nociceptors where they become sensitized and increase patient pain.

## Introduction

Endometriosis is a common disease characterized by chronic pelvic pain^1–6^ which impacts approximately 70% of patients, 37% of which report severe symptoms^7,8^. The pain is resistant to many analgesics including opioids and anti-inflammatory agents^9–12^. Hormonal therapies are often implemented and reduce endometriosis-associated pain in a subset of patients^13,14^. However, 8-40% of patients do not respond to hormonal therapies, and these therapies are often stopped due to side effects^9,13^, raising the need for new treatment options with improved efficacy and reduced side effect burden. The disease is defined by ectopic growth of endometrial-like tissue forming lesions within the abdominal cavity, most commonly on the peritoneal wall^7,20–22^. Laparoscopic surgery to identify and resect lesions reduces pain-related symptoms in about 65% of patients. However, lesions frequently recur (28% at 18 months, 40% at 9 nine years), necessitating repeat procedures^9,15,16^. The chronic nature of endometriosis-associated pelvic pain leads to healthcare costs of 69.4 billion US dollars/year. The high impact on the patients’ quality of life highlights the need for better understanding of the mechanisms underlying endometriosis to improve treatments, particularly those that target the pain of endometriosis^6,17,18^.

Endometriosis is a complex inflammatory, and endocrine-dependent disease composed of multiple interacting cellular compartments, including ectopic endometrial cells (gland-associated epithelial and stromal cells), immune populations, vasculature, and innervating axons from the dorsal root ganglia (DRG) and sympathetic ganglia^19–23^. Histologically, lesions are defined by endometrial glands containing an epithelial layer surrounded by a dense stromal cell layer, resembling glands in the eutopic endometrium^24^. While substantial research has focused on hormonal and immune contributions to disease development, mechanisms underlying the patient pain experience remain poorly defined^25–30^. Clinical endometriosis staging describes the severity of the disease based on lesion number, depth, and anatomical distribution^31^. Despite clear presentation differences between patients, disease stage does not reliably predict pain severity^32–34^. This disconnect suggests that additional factors contribute to the pain of endometriosis.

DRG sensory neurons include nociceptors that transmit noxious peripheral signals to the central nervous system, including endocrine and inflammatory signals in tissue^35–37^. Clinical studies suggest that endometrial lesions are hyper-innervated relative to normal peritoneum^22,38,39^. Critically, the density of innervation correlates to patient pain^39–41^ and is reduced by hormonal treatment used to manage the pain^42^. Surgical resection of lesions reduces pain transiently before lesions recur^13,43,44^, linking symptoms to innervated ectopic tissue. Further, ablation of the uterosacral-nerve can reduce patient pain^45^. However, these observations remain largely correlative and do not functionally assess the neurons that innervate lesions since these tissues are not accessible in the clinic. As a result, the excitability, molecular identity, and recruitment mechanisms of lesion-innervating neurons remain poorly defined. Recent advances targeting DRG neurons to reduce pain in other conditions highlight the importance of characterizing the sensory neuron component of this disease^46^.

To address current limitations, preclinical mouse models of endometriosis have been developed to enable mechanistic investigation of the disease^47–49^. Although mice do not menstruate, introduction of endometrial tissue into the peritoneal cavity results in the formation of ectopic lesions with key features of the human disease, including implantation site and histological similarities^47^. In these models, ablation of nociceptors reduces pain-like behaviors, supporting a functional role for sensory neurons in endometriosis-associated pain^30^. Given that lesions are ectopic tissues that become innervated during or after formation, it is possible that processes that drive DRG sensory neuron innervation could contribute to endometriosis pain, and possibly the disease itself^30^. Innervation of peripheral tissues is a regulated process in which axons are guided by local molecular cues^50–52^. During development, target-released glial cell line-derived neurotrophic factor (GDNF) promotes the growth of sensory neurons that express the Ret receptor tyrosine kinase and the GDNF co-receptor GFRα1^53–57^. Further, in the context of injury, GDNF/GFRα1/Ret signaling in nociceptors contributes to pain-like behaviors^58,59^. Given these dual roles in target innervation and modulation of neuronal excitability, it is possible that this growth factor system is mechanistically related to disease progression and pain severity in endometriosis. In conjunction with the previously described presence of GDNF and other growth factors in endometriosis^60,61^, this signaling system is poised to be disease modifying.

In this study, we investigate the DRG neurons which innervate endometriosis lesions. In the mouse model, we find that lesion-innervating DRG neurons are small-diameter and bind IB4, a marker for the GDNF Family Ligand (GFL) receptor Ret, consistent with their identification as nociceptors. We show that these neurons are hyperexcitable compared to neurons innervating the adjacent peritoneal wall. In patient lesions, GDNF is expressed in the stromal layer of endometrial glands nearby innervating axons that are frequently GFRα1-positive. Finally, we find that the density of innervation and abundance of GDNF and GFRα1 positively correlate with patient-reported pain. These data suggest novel, non-hormonal targets for the management of endometriosis pain.

## Results

### Induction of endometriosis in a mouse model drives chronic abdominal pain-like behaviors

To investigate mechanistic changes due to endometriosis, we first adopted a preclinical mouse model^47^. In this validated model, eutopic endometrium of a donor animal is injected into the peritoneal cavity of a host animal (Endo). Littermate control animals were injected with media without tissue (Sham). The disease model was allowed to develop over eight weeks (**Fig. 1A**) which resulted in large, vesicle-like lesions that frequently formed on the ventral peritoneal wall, similar to human vesicular endometrial-like lesions^31,62–64^ (**Fig. 1B**). Histologic analysis of mouse and human lesions demonstrate that samples from both species present with endometrial-like glands (*G*) enveloped by a dense stromal cell compartment (*Str.*), immediately adjacent to the gland-epithelial layer. Inflammation/fibrosis (*Infl.*) is also present at the border of the gland-related stroma and in nearby tissue, as expected^60^ (**Fig. 1B**). In the animal model, we found during careful gross postmortem dissection that ∼78.6% of Endo animals formed visually detectable lesions at an average of ∼2 lesions/animal. (**Fig. 1C**). Endo animals were apparently healthy and indistinguishable from Shams with no difference in weight over the development of the model (**Fig. 1D**). Behavioral assessments at five weeks and eight weeks post induction revealed abdominal mechanical hypersensitivity of Endo animals compared to Shams (**Fig. 1E-F**), as previously reported^47^. Spontaneous pain-like behaviors also emerged in Endo animals including writhing^65^ and “abdominal squashing” characterized by pressing of the abdomen against the floor grate^47^ (**Fig. 1G-H**). No difference in grooming/licking directed to the abdomen was found between groups (**Fig. 1I**). Given the clinical prevalence of dysmenorrhea^16^, we grouped pain-like behaviors across the mouse estrous cycle determined by visual inspection of the external genitalia^66^. We found no effects of estrous phase on mechanical sensitivity but observed an increase in the number of “abdominal squashing” bouts in Endo animals in metestrus relative to those in diestrus at eight weeks post transfer. No behavioral differences were detected across estrous phases in Sham animals, although we are not sufficiently powered to detect changes across all phases (**Supp. Fig. 1**). Together, these data indicate that the mouse endometriosis model recapitulates key aspects of the human pain phenotype, including spontaneous behaviors, and might therefore provide insights into the human disease.

**Figure 1.**
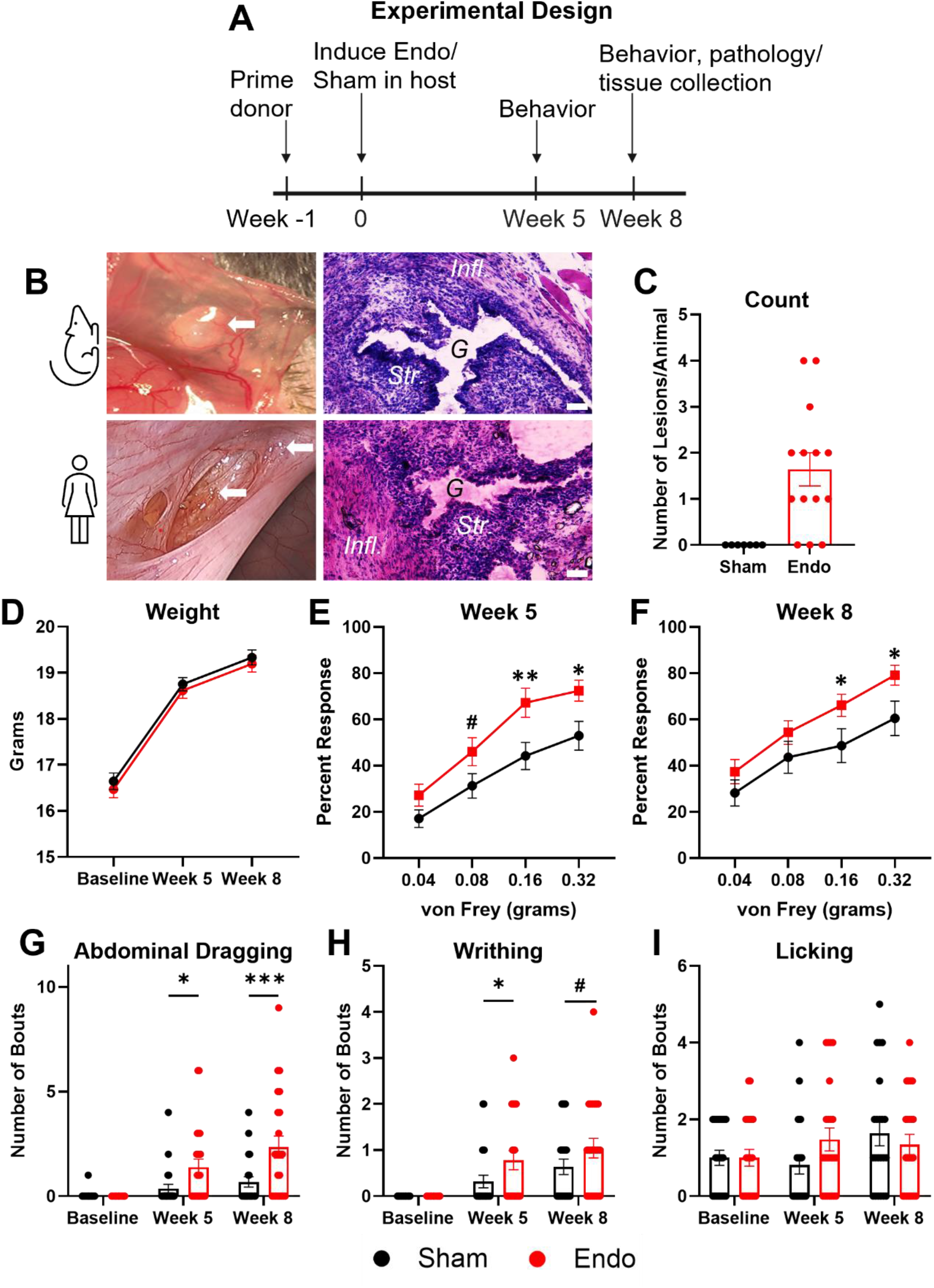
Induction and validation of a mouse model of endometriosis. **A.** Schematic of the experimental design used to induce and test the model of endometriosis (Endo) or control (Sham) in mice. **B.** Gross anatomy representation and histological analyses of human vesicular endometriosis and mouse lesions appear similar; both present with a gland (marked by a “*G*”) surrounded by endometrial stroma (marked by “*Str.*”) and tissue inflammation (marked by “*Infl.*”). **C.** In Endo animals, lesions are detected at an average rate of 1.6/animal. **D.** Average weight of animals between groups is not different at measured times. **E.** Endo and littermate Sham animals were tested for mechanical sensitivity of their abdomen five weeks following the induction of the model. Percent withdrawal to calibrated von Frey fibers were recorded in response to 10 abdominal applications. Endo animals responded more frequently with robust withdrawal behaviors to 0.16 and 0.32 gram-force (Two way RM ANOVA, Tukey’s; *p=0.012, **p=0.003, #p=0.056; n=24-25 animals/group). **F.** Repeated behavior at eight weeks following model induction replicates these results (Two way RM ANOVA, Tukey’s; *p<0.05; n=22-23 animals/group). **G.** Spontaneous pain-like behaviors were also observed for 10 minutes. Abdominal dragging, defined by the animal pressing its abdomen against the grate it stands on, was increased in Endo animals relative to Shams at five and eight weeks post induction (Two way RM ANOVA, Tukey’s; *p=0.017, ***p=0.0001; n=22-23 animals/group). **H.** Writhing behavior was also increased in animals at five weeks, although not statistically different than Shams at eight weeks (Two way RM ANOVA, Tukey’s; *p=0.032, #p=0.06; n=22-23 animals/group). **I.** No differences between groups was noted in abdominal-directed licking behaviors at any time point (Two way RM ANOVA, Tukey’s; n=22-23 animals/group). Scale=50 µm. Data represented as mean +/- SEM.

### Electrophysiological properties of DRG sensory neurons innervating peritoneal lesions in mouse

Given the clinical data suggesting that innervation density in lesions is related to patient-reported pain^39–41^, and because the lesions contain inflammatory and hormonal factors that can sensitize neurons^67^, we sought to functionally characterize lesion-innervating neurons. At the conclusion of behavioral testing, we retrogradely labeled DRG neurons that innervate lesions by micro-injecting WGA into peritoneal lesions (Lesion). As a control, we injected WGA into the wall of the peritoneum where lesions were absent (Endo Wall; **Fig. 2A-B**). In addition, we injected WGA into the peritoneal wall of littermate sham animals at a location where lesions often form (Sham Wall). Animals were allowed to recover for three to seven days to allow sufficient time for retrograde transport of the dye to the DRG cell bodies of neurons innervating the injection site. WGA injected into lesions remains in lesions without spreading to adjacent non-lesion peritoneum (**Supp. Fig. 2**). Concurrently, we dissected and dissociated DRGs for *in vitro* characterization of neuronal excitability by patch clamp electrophysiology of WGA-labeled neurons (**Fig. 2C**). We restricted electrophysiology analyses to small-diameter neurons (≤30 µm, putative nociceptors^68–70^; **Fig. 2D**). We found that the resting membrane potential (RMP) of Lesion-innervating neurons was significantly more depolarized at rest compared to neurons innervating the Sham Wall (**Fig. 2E**). Interestingly, Endo Wall-innervating neurons showed a trend toward being depolarized relative to Sham Wall-innervating neurons, although the differences were not statistically significant (**Fig. 2E**). Among the lesion-innervating neurons, we observed several instances of spontaneous firing at rest, while this was never seen in the controls (**Fig. 2F**). We found no difference in the incidence of single/repetitive firing of Lesion-innervating neurons compared to control neurons (**Fig. 2G**). For experiments evaluating depolarization-evoked excitability, all neurons were held at -60 mV. Consecutive step injections revealed that the rheobase (minimum amount of current necessary to fire an action potential (AP)) of Lesion-innervating neurons was significantly lower compared to control neurons (**Fig. 2H**, representative image **Fig. 2H′**). Evaluation of AP dynamics revealed that the peak amplitude of the AP was significantly reduced in Lesion-innervating neurons relative to control neurons (**Fig. 2I**, representative image **Fig. 2I′**). Additional electrophysiological properties were also tested with no significant differences in Lesion-innervating neurons compared to Sham Wall-innervating neurons (**Suppl. Fig. 2**). These results suggest that Lesion-innervating neurons are more excitable and better poised to readily fire APs relative to neurons that innervate the peritoneal wall.

**Figure 2.**
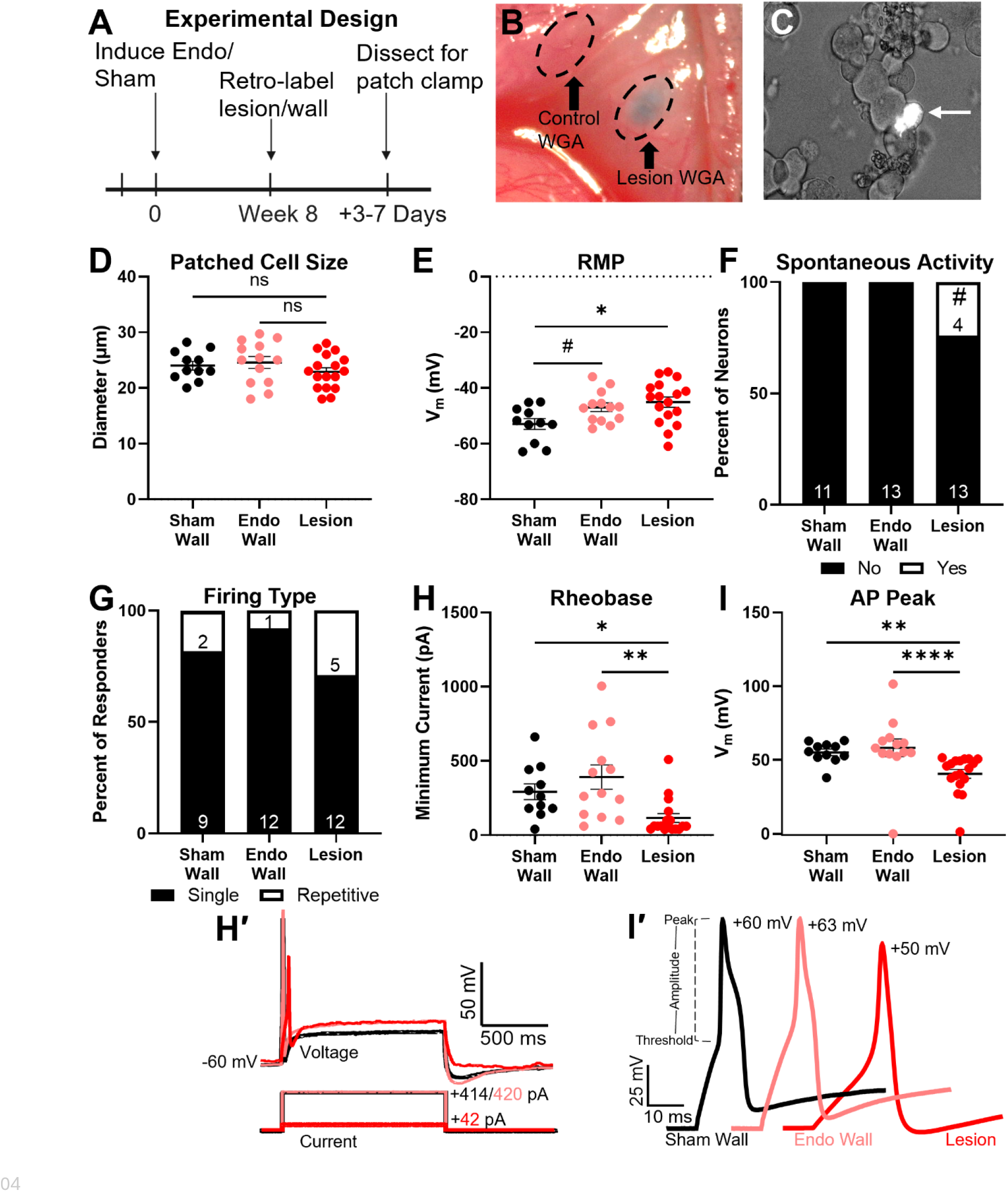
Lesion innervating neurons are more excitable compared to size-matched controls. **A.** Schematic of the experimental design depicting the model generation and time of laparotomy and injection of retrograde WGA into either **(B)** the ventral peritoneal wall or a peritoneal lesion. **C.** Representative image of neurons *in vitro* with one labeled by WGA retrograde dye (arrow). **D.** Neurons that were ≤30 µm were selected for analysis with no difference in size between groups (One way ANOVA, Tukey’s). **E.** The resting membrane potential (RMP) of lesion innervating neurons is significantly depolarized relative to Sham Wall innervating neurons. Endo Wall innervating neurons are trending toward more depolarized potentials compared to Sham wall (One way ANOVA, Tukey’s; *p=0.013, #p=0.09). **F.** 23.5% of lesion-innervating neurons fire APs at rest while Sham Wall-and Endo Wall-innervating neurons do not fire APs at rest (Fisher’s exact; #p=0.055; number of cells indicated within the bars). **G.** The proportion of neurons that fire multiple APs or single APs is not different between groups (Fisher’s exact; p=0.32; number of cells indicated within the bars). **H and H′.** An AP was evoked from lesion-innervating neurons on average at lower step current injected (rheobase) compared to both controls (Kruskal-Wallis, Dunn’s; *p=0.025, **p=0.0031). **I and I′.** The average peak of APs in lesion-innervating neurons was significantly lower compared to wall-innervating neurons (Kruskal-Wallis, Dunn’s; **p=0.0022, ****p<0.0001). n=11 Sham Wall-innervating neurons sampled from 3 animals, 13 Endo Wall-innervating neurons sampled from 4 animals, 17 Lesion-innervating neurons from 7 animals. Data represented as mean +/- SEM or percentage bars.

During recordings it became apparent that the number of small-diameter neurons in the Lesion-innervating group was greater than the number of small-diameter neurons in the peritoneal wall-innervating groups. While electrophysiological properties were all measured in small-diameter neurons (**Fig. 2**), analysis of all WGA-labeled neurons in culture revealed an overall smaller average diameter of Lesion-innervating neurons compared to control neurons (**Fig. 3A**). Further, the proportion of all neurons in culture that were small-diameter (≤30 µm) among Lesion-innervating neurons was significantly greater compared to peritoneal wall-innervating groups (**Fig. 3B**). Given these results, we hypothesized that hyperexcitable lesion-innervating neurons represent a distinct subpopulation of putative nociceptors.

**Figure 3.**
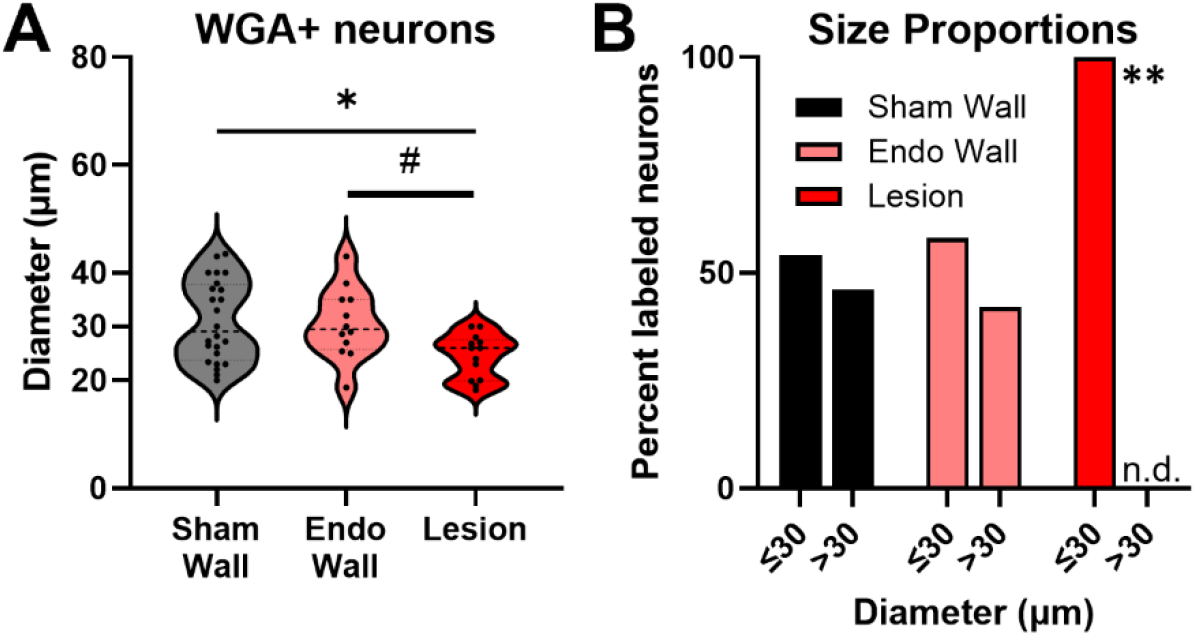
Neurons which innervate lesions are small-diameter *in vitro*. **A.** Coverslips of DRG neurons innervating Sham Wall, Endo Wall, or Lesions were scanned for all WGA+ cells. Lesion-innervating neurons are significantly smaller than Sham Wall-innervating neurons, and trending smaller compared to Endo Wall-innervating neurons (One way ANOVA, Tukey’s; #p=0.06, *p=0.015). **B.** Small-diameter (≤30 µm) neurons are more abundant than large diameter (˃30 µm) lesion-innervating neurons, but other groups have equal WGA+ representation of small and large diameter neurons (Fisher’s exact; **p=0.0061). n=24 Sham Wall, 12 Endo Wall, 13 Lesion. Data represented as the median and quartiles.

### Lesions are innervated by small-diameter, IB4/Ret positive neurons

To determine the identity of WGA labeled neurons, we performed immunohistochemistry on DRG tissue from Sham Wall (**Fig. 4A-A′′**) and Lesion (**Fig. 4B-B′′**) traced animals. We found that micro-injections into these locations resulted in WGA labeling in lower thoracic (T11-13) and upper lumbar (L1-2) DRG levels, consistent with previously mapped innervation patterns^71,72^. Isolectin B4 (IB4)-binding is classically used to identify a subset of DRG neurons that are small-diameter and Ret positive, indicating responsiveness to GDNF family ligands^54,59,73^. Considering our *in vitro* evidence that lesion-innervating neurons are small-diameter (**Fig. 3**), we tested whether DRG neurons retrogradely labeled from lesions also bound IB4. The proportion of WGA-labeled neurons did not differ between groups (**Fig. 4C**). However, the average diameter of Lesion-innervating neurons was significantly smaller than the average diameter of Sham Wall-innervating neurons (**Fig. 4D**). A distribution analysis of diameters binned by size demonstrates that Lesion-innervating neurons skew towards small-diameter, putative nociceptors, whereas Sham Wall-innervating neurons have a more widespread size distribution (**Fig. 4E**). Finally, consistent with the differences in sizes, quantification of IB4-positive neurons reveals that Lesion-innervating neurons bind IB4 significantly more frequently than Sham Wall-innervating neurons (**Fig. 4F**). Together, these data support the *in vitro* observation that Lesion-innervating DRG neurons are predominantly small-diameter neurons belonging to the Ret expressing subpopulation of nociceptors and suggest that GFL receptors (GFRs) and their associated ligands may play a role in the innervation of endometriosis lesions.

**Figure 4.**
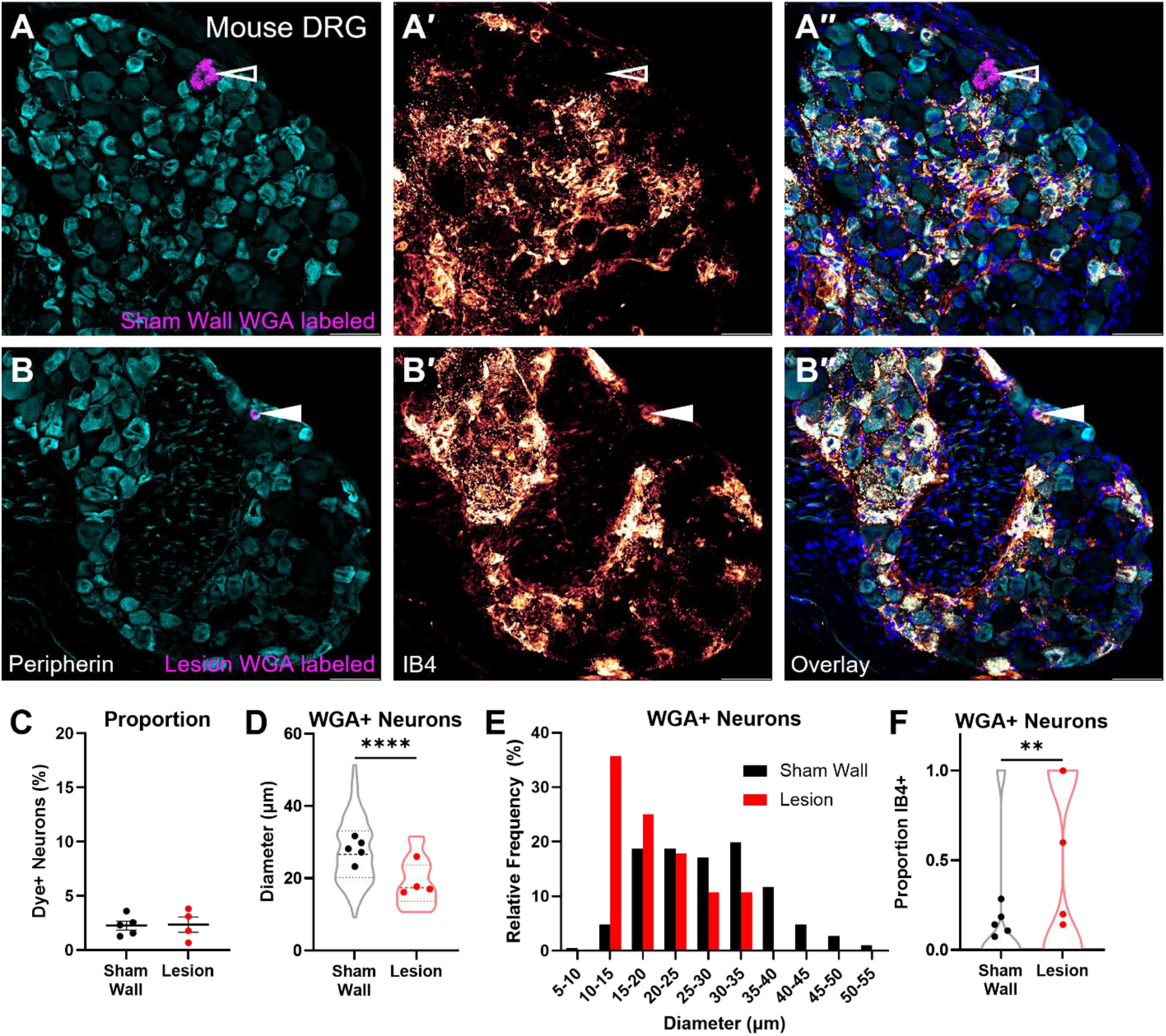
DRG neurons that innervate peritoneal lesions are preferentially small-diameter and IB4+. **A.** Representative images of a Sham Wall-innervating neuron (peripherin; teal) labeled with retrograde dye (WGA; magenta) but not co-stained with IB4 (orange/yellow; open arrow). **B.** Representative image of a Lesion-innervating neuron that co-stains with IB4 (closed arrow). **C.** The average proportion of WGA-labeled neurons relative to Peripherin+ neurons in WGA-present sections is unchanged by innervation target (Student’s t test). **D.** The diameter of Lesion-innervating neurons is smaller than Sham Wall-innervating neurons (Mann-Whitney U test; ****p<0.0001). **E.** Size-Frequency histograms reveals a greater frequency of small-diameter neurons in DRG retrogradely labeled from lesions compared with DRG labeled from the peritoneal wall of Sham mice. **F.** Co-staining of WGA marked neurons with IB4 occurs more frequently in Lesion-innervating neurons relative to Sham Wall-innervating neurons (Mann-Whitney U test, **p=0.0026). Data acquired from DRG segments T11-L2; Sham Wall n=187 neurons, 5 animals; Lesion n=28 neurons, 5 animals with no dye detected in one animal. Data represented as mean +/- SEM or mean and interquartile range of WGA+ neurons with averages per animal indicated by individual dots. Scale=50 µm.

We and others have previously shown, and confirm here using a Ret-reporter mouse line^74^, that IB4-labeled neurons almost exclusively also express Ret, particularly in the small-diameter population (**Supp. Fig. 3**)^54,75^. To determine if the Ret+ population of small-diameter DRG neurons are functionally distinct from Ret-small-diameter neurons in naïve animals, we performed patch clamp electrophysiology of small-diameter DRG neurons in the Ret-reporter mouse. We found that Ret+ neurons have a higher rheobase (less excitable) than Ret-neurons. Action potential waveform analysis found that Ret+ neurons have a lower amplitude and peak compared to Ret-neurons (**Supp. Fig. 3**). Together, these data demonstrate that although small-diameter Ret+ DRG neurons are inherently less excitable than Ret-DRG neurons, when this population innervates lesions they show increased excitability compared to small-diameter neurons that innervate the peritoneal wall and are often Ret-. Overall, our data is consistent with a potential role for GFL/Ret signaling in lesion innervation that might contribute to sensitization and pain in this endometriosis mouse model.

### Endometriosis patient-reported pain is correlated with stromal GDNF levels and neuronal GFRα1 levels

To explore the potential role of GDNF/Ret signaling in clinical endometriosis, we recruited patients scheduled for minimally invasive gynecological surgery to diagnose and resect endometriosis lesions. During a study visit prior to scheduled surgery, patients completed the Endometriosis Health Profile-30 Pain Scale (EHP-30) and reported their endometriosis-related pelvic pain intensity over the last 30 days using a scale ranging from 0 (no pain) to 10 (worst pain imaginable). Fourteen participants who had a self-reported dynamic range of pain and confirmed endometriosis lesions that were removed during the surgery were included in this study. Consistent with some prior reports, clinically assigned disease stage was not related to patient reported pain (**Table 1**)^32–34^.

**Table 1.** Patient demographics, clinical characteristics, and pain scores. Data include self-reported endometriosis pain intensity, calculated EHP-30 pain scores, and AAGL surgical stage. Anatomical location and corresponding dataset for each sample (proteomics and/or IHC) are also noted. N/A staging indicates that stage was not assigned.

| Pain Intensity | EHP-30 Score | Age | Race | Ethnicity | Lesion location | AAGL Stage | Proteomics sample | IHC sample |
| --- | --- | --- | --- | --- | --- | --- | --- | --- |
| 2 | 43.1 | 25 | Caucasian | Non-Hispanic | Pelvic side wall | N/A | No | Yes |
| 3 | 29.5 | 28 | Caucasian | Non-Hispanic | Pelvic side wall | 1 | Yes | Yes |
| 4 | 70.5 | 38 | Caucasian | Non-Hispanic | Pelvic side wall | 2 | Yes | No |
| 4 | 38.6 | 30 | Caucasian | Non-Hispanic | Pelvic brim | 4 | No | Yes |
| 5 | 20.5 | 28 | Caucasian | Non-Hispanic | Perirectal | 3 | Yes | No |
| 5 | 18.2 | 45 | Caucasian | Non-Hispanic | Pelvic side wall | 2 | Yes | No |
| 6 | 36.4 | 37 | Bi-racial | Hispanic | Peritoneum | 3 | Yes | No |
| 6 | 68.1 | 21 | Caucasian | Non-Hispanic | Posterior Cul de Sac | N/A | Yes | No |
| 7 | 56.8 | 29 | Caucasian | Non-Hispanic | Pelvic floor | 4 | Yes | No |
| 7 | 54.5 | 19 | Caucasian | Non-Hispanic | Ovarian fossa | N/A | Yes | No |
| 7 | 59.0 | 32 | Caucasian | Non-Hispanic | Side wall | 1 | Yes | Yes |
| 8 | 72.3 | 25 | Caucasian | Non-Hispanic | Rectovaginal pelvic floor | 4 | Yes | Yes |
| 9 | 63.6 | 28 | Caucasian | Non-Hispanic | Uterosacral ligament | 3 | Yes | No |
| 9 | 65.9 | 34 | Caucasian | Non-Hispanic | Posterior Cul de Sac | 1 | Yes | Yes |

We sought to identify molecular markers that correlate with patient pain. We first compared lesions from patients who reported high/severe pelvic pain (pain ratings = 7-9; N=6) to lesions from patients reporting mild/moderate pain (pain ratings = 3-6; N=6) by performing unbiased bulk proteomics assaying >5,000 proteins. We found multiple growth factors enriched in lesions from high/severe pain participants including GDNF, NOTCH1, ARTN, NT3, and BDNF (**Fig. 5A**). Although these differences are not statistically significant, a principal component analysis reveals a cluster of high-pain samples with high expression of GDNF (**Supp. Fig. 4**). Interestingly, binning patients into groups of mild (pain intensity = 3-4; N=2), moderate (intensity = 5-6; N=4), high (intensity = 7; N=3), and severe pain (intensity = 8-9; N=3) reveals a clear trend where GDNF levels are positively associated with clinical pain scores (**Fig. 5B**). Analysis of publicly available endometriosis lesion datasets indicates that GDNF is likely expressed in stromal cells within endometriosis lesions although smooth muscle cells and B cells also express this gene^76^. We leveraged our proteomic dataset to assess relationships between GDNF and stromal cell markers (COL4A1, CD10, ESR1 and PGR), some of which are used clinically to diagnose endometriosis^77,78^. We found that the levels of most stromal factors are tightly associated with GDNF, but B cell and smooth muscle cell markers do not correlate with GDNF levels (**Supp. Fig. 4**). Normalization of GDNF levels to COL4A1 (pan-stromal cell marker, **Fig. 5C**) as well as CD10 (diagnostic endometriosis stroma marker; **Fig. 5D**) suggests that the association between pelvic pain and GDNF levels might be due to stromal cell expression of GDNF. Further, we found by immunohistochemistry that GDNF is restricted to endometrial glands, most commonly in the stromal cell compartment of patient lesions (**Fig. 5E**) and of the mouse model lesions (**Supp. Fig. 5**). Finally, pathway enrichment analysis implicates both stromal cell expansion and inflammatory pathways to be enriched in lesions from patients reporting high/severe pain (**Supp. Fig. 4**). Together, these data support a possible relationship between lesion GDNF and endometriosis pain and strongly associate GDNF with the lesion stromal cell compartment.

**Figure 5.**
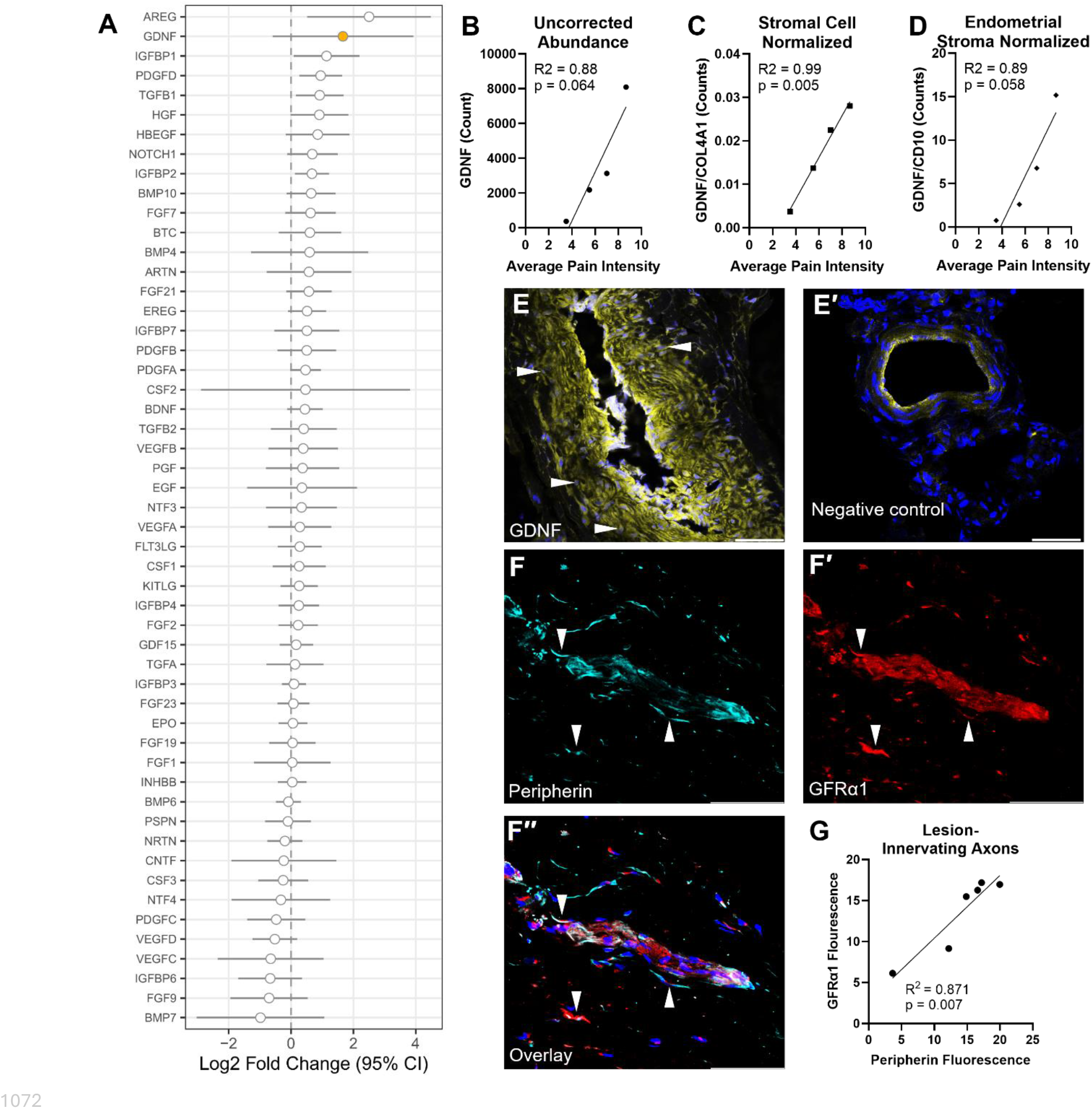
Identification, abundance and probable cellular compartments of GDNF expression in lesions from patients with pain-defined endometriosis. **A.** Forest plot of proteomics data demonstrates the magnitude of change (Log2 Fold Change) of growth factors in endometriosis patients with high/severe endometriosis-related pelvic pain (Intensity = 7-9; N=6) compared to patients with mild/moderate pain (Intensity = 3-6; N=6). A right-shift is indicative of enrichment in the high/severe pain group. **B.** Binned proteomic counts by mild (Intensity = 3-4; N=2), moderate (Intensity = 5-6; N=4), high (Intensity = 7; N=3), and severe (Intensity = 8-9; N=3) reveals a trending positive association of GDNF with pain intensity. **C.** Normalization of GDNF counts to total stromal cell (COL4A1) counts or (**D**) specifically stroma associated with endometrial glands (CD10) counts, indicates strong associations with average reported pain intensity for each. **E.** GDNF (yellow) expression is restricted to the stromal cell layer of endometriosis glands and is absent in sections which were not exposed to the primary antibody (**E′**). Arrows indicate GDNF positive stroma. **F-F′′.** Immunohistochemical representative images and (**G**) quantification of Peripherin and GFRα1 demonstrate a strong correlation in their expression levels. Arrowheads indicate double positive axons. Simple linear regressions, goodness of fit and significance values indicated on individual panels. Scale=50 µm.

Publicly available human DRG datasets confirm that GFRα1 is highly expressed in subsets of neurons consistent with nociceptors^79^. In our proteomic dataset, we found that GFRα1 expression, but not GFRα2/3, correlates with the abundance of Peripherin, a marker for the peripheral nervous system including both sensory and sympathetic axons (**Supp. Fig. 4**). Immunohistochemical analysis confirmed a strong correlation between GFRα1 and Peripherin in axons innervating endometrial-associated stroma and glands (**Fig. 5F-G**). Associations in the proteomics dataset reveal that GFRα1 levels correlate with TRPV1 levels (sensory axon specific) but not TH levels (sympathetic axon specific)^80,81^. This suggests that the GFRα1+ axons innervating lesions are sensory neurons (**Supp. Fig. 4**) which is consistent with prior reports that demonstrate a reduction in sympathetic axons in lesions compared to healthy peritoneal tissue^82^. These data, combined with the expression of GDNF in lesions, inspired further testing and validation to determine if a GDNF-GFRα1 axis is modified in lesions based on pain score.

### Patient endometriosis pain is related to levels of GDNF-GFRα1 in lesions

After identifying the putative sources of GDNF and GFRα1 in lesions, we next sought to validate whether expression of GDNF and GFRα1 in endometriosis lesions is related to clinical pain scores. Given the apparent containment of GDNF to the stromal layer of endometrial glands, we restricted our histological quantification to endometrial glands. We found a strong trend between cell density-normalized GDNF expression level mean fluorescence intensity (MFI) and clinical pain, consistent with the proteomic finding that GDNF levels are related to pain intensity (**Fig. 6A-C**), though this was not statistically significant. Importantly, the size of the gland and the density of cells in the region analyzed did not correlate with pain (**Supp. Table 1**). Analysis of Peripherin in endometrial glands supports prior reports that patient pain is related to lesion axonal density^39–41^ (**Fig. 6D-F**), here localized specifically to glands. Robust innervation density is also found in the lesions dissected from the animal model, demonstrating a similar pattern of innervation as human (**Supp. Fig. 5**). Finally, we quantified the amount of GFRα1 fluorescence and found that GFRα1 also displays a strong trend to be correlated with patient reported pain (**Fig. 6G-I**). GDNF, Peripherin and GFRα1 MFI were converted to a composite z-score to analyze GDNF-GFRα1 signaling in our dataset^83^. This score was significantly correlated to clinical pain (**Fig. 6J**) and displayed a positive trend to associate with patient’s EHP-30 pain score (**Fig. 6K**). Together, these data provide correlative evidence in human samples indicating that stromal GDNF and axonal GFRα1 expression levels are related to clinical pain intensity supporting the mechanistic findings in the animal model.

**Figure 6.**
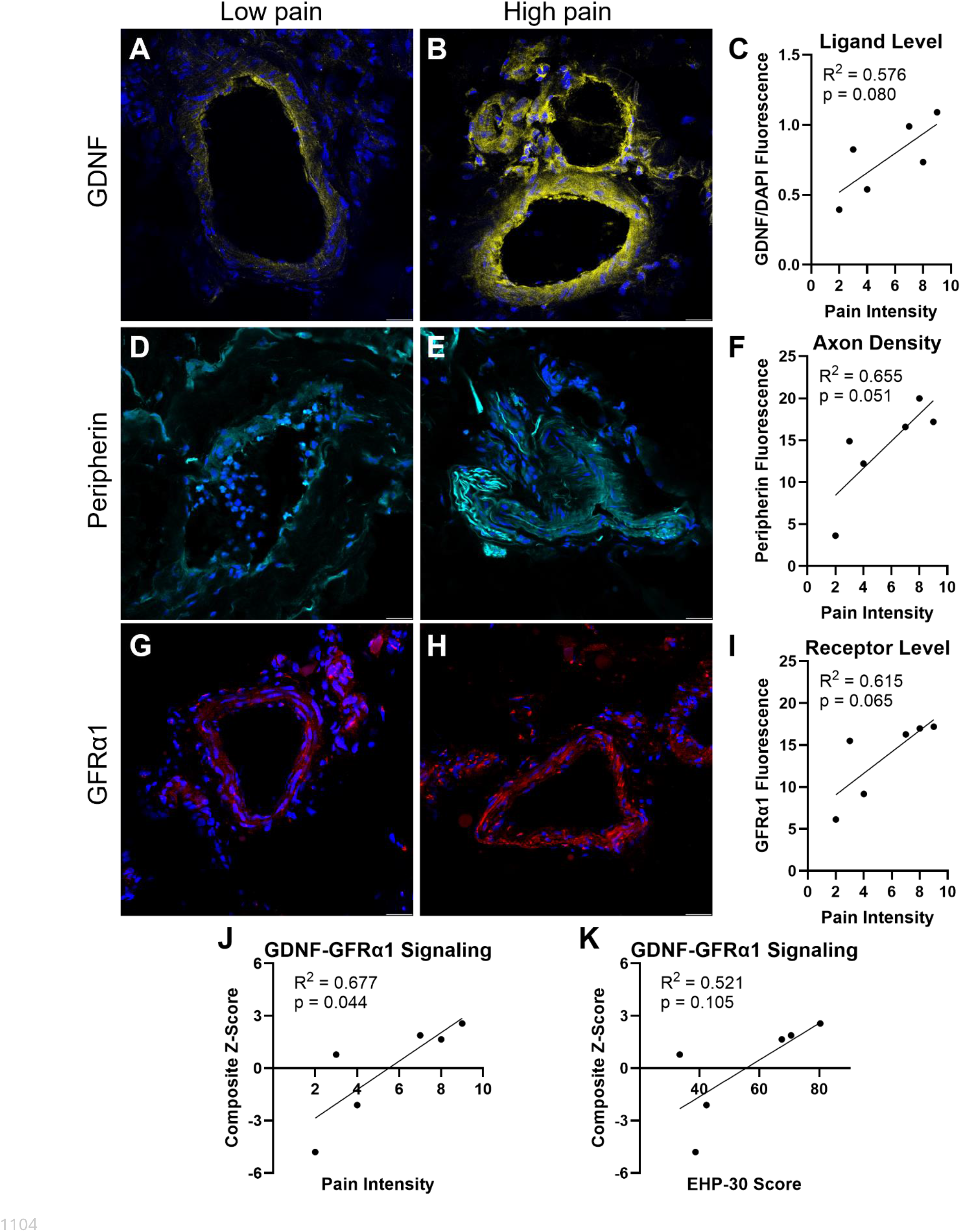
Associations between GDNF and GFRα1 expression levels in lesions with patient-reported pain. A-C. Representative images of GDNF in low and high pain reporting patients and quantification of mean fluorescence intensity of GDNF/DAPI versus reported pain intensity. **D-E.** Visualization of axons by peripherin staining at endometrial glands and quantification of levels by mean fluorescence intensity versus patient reported pain. **G-I.** Images demonstrating GFRα1 levels at endometrial glands and quantification by mean fluorescence intensity plotted against patient reported pain. **J-K.** MFI was converted to z-scores to make a composite of GDNF/DAPI, Peripherin, and GFRα1 to associate GDNF-GFRα1 signaling potential at endometrial glands in patients. Composite z-scores are correlated with reported pain intensity (**J**) and demonstrate a trend with calculated EHP-30 pain score (**K**). Simple linear regressions, N=6 (averaged across all glands present in at least 3 non-consecutive sections), goodness of fit and significance values indicated on individual panels. Scale=50 µm.

## Discussion

We report data suggesting distinct properties and identities of DRG neurons that innervate endometrial-like lesions in both a mouse model of the disease and pain-phenotyped participants who have endometriosis (**Fig. 7**). We confirm that the mouse model recapitulates prominent features of the pain experienced by humans including spontaneous, pelvic-directed pain-like behaviors (**Fig. 1**)^47^. We demonstrate, for the first time, that mouse endometrial-like lesions are innervated by DRG neurons that are hyperexcitable compared to control neurons (**Fig. 2**) and which we identified to be the Ret/IB4 subpopulation of small-diameter nociceptors (**Figs. 3-4**)^54^. These findings in the mouse model prompted us to examine the innervation of endometrial lesions from human patients who self-reported their pelvic pain. We identified GFL signaling components in human lesions (**Fig. 5**) and revealed that levels of GDNF, and its co-receptor GFRα1 on lesion-innervating axons, were predictive of patient-reported pain (**Fig. 6**). These data are consistent with the hypothesis that GDNF expressed in lesions drives neuronal cell-type specific innervation of lesions followed by neuronal sensitization, increased excitability and pelvic pain.

**Figure 7.**
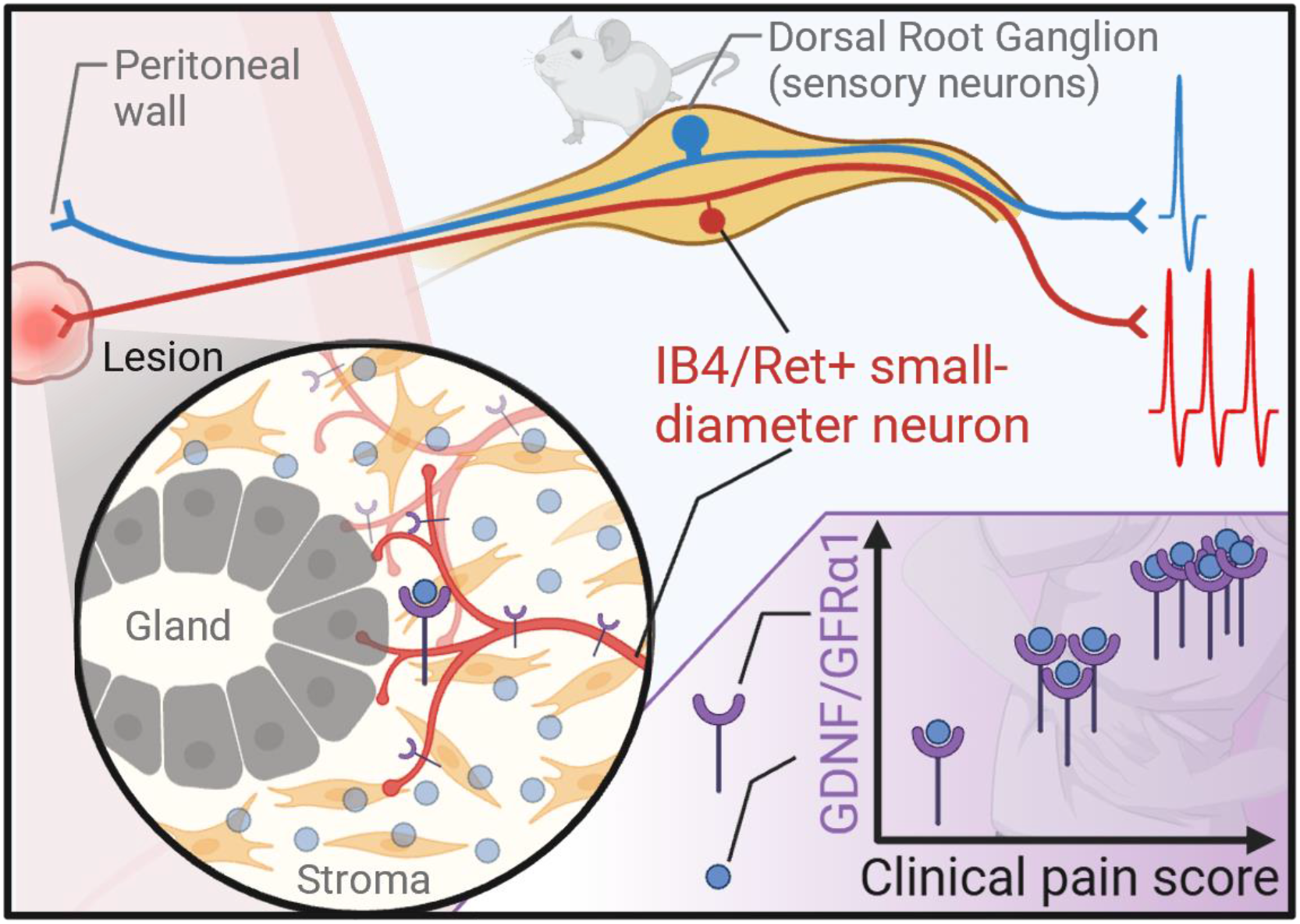
Schematic summary of the results. Dorsal root ganglion sensory neurons which innervate lesions are smaller in diameter than those which innervate control peritoneal wall. They are also preferentially IB4/Ret+ and hyperexcitable. The same subtype of sensory neurons (GFRα1+) innervate patient lesions most densely in the stromal cell compartment where GDNF is present. In participants which self-reported their endometriosis-associated pain, the levels of GDNF, innervation and GFRα1 associate with their pain score. Graphic made using BioRender.

Human endometriosis lesions consist of various tissues/cell types including implantation site tissue (peritoneum), endometrial-like tissue (epithelial cells, stromal cells), vascular tissues, immune cells, and innervating axons^19,20^. The molecular landscape in endometrial-like lesions is consequently complex^20,27–29,84–86^. We reasoned that evaluating the proteome of lesions from patients who reported a dynamic range of pelvic pain might reveal novel therapeutic strategies targeting pain. Our data support GDNF as a strong candidate that may link patient pain and lesion biology to neuronal recruitment. It is well established that GDNF-GFRα1 signaling regulates the targeting and survival of specific sensory neuron populations during development^54–56^. It is therefore possible that similar mechanisms are reactivated in endometriosis, similar to neurotrophic factor-mediated targeting of axons to tumors in certain cancers^87,88^. According to publicly available human datasets, GDNF is expressed at low levels in the stromal cell compartment of the eutopic endometrium in the uterus (putative source of endometrial cells that form lesions)^76^. It is therefore possible that after/during ectopic attachment of endometrial cells to the peritoneal wall, GDNF is upregulated^60^ (**Fig. 5**) and capable of increasing GFRα1-positive sensory neuron innervation (**Supp. Fig. 5**). The identification of this neuronal subtype in human lesions is consistent with the mouse data, which revealed that DRG neurons innervating lesions are often small-diameter and IB4-positive, designating them as Ret+ nociceptors (**Fig. 4 and Supp. Fig. 3**). Functional experiments to reduce GDNF in lesion-fated stromal cells would therefore be a valuable tool to determine the requirement for GDNF to drive pain-like behaviors, innervation, and lesion formation, and would provide further rationale for a therapeutic approach targeting GFL/GFRα1/Ret signaling.

The prevalence of this growth factor signaling axis in endometrial lesions in mouse endometrial-like lesions prompted us to examine the relationship between GFRα1 and GDNF expression in human endometriosis lesions and clinical pain (**Fig. 6**). In human endometriosis lesions, we find that the level of GDNF, putatively expressed by stromal cells of the endometrial gland, and the level of the GFRα1 expressed by DRG neurons that innervate lesions, are positively correlated with clinical pain intensity and show a positive correlation trend with endometriosis health profile. These data support further examination of the role of the GDNF/GFRα1/Ret signaling pathway in endometriosis pain. Together, these data support the hypothesis that the level of GFLs in lesions impact patient pain by increasing recruitment of nociceptors to lesions.

In addition to GDNF, we also identified several other neurotrophic factors that are non-significantly enriched in lesions from patients who reported more pain. BDNF and NT3 are notable due to their roles in endometriosis pathogenesis^89,90^ while ARTN is of interest because it preferentially binds to the GFL co-receptor GFRα3 which is expressed on a different subset of DRG sensory neurons than GFRα1^53,56^ (**Fig. 5**). It is possible that ARTN/GFRα3 signaling represents a parallel pathway to GDNF/GFRα1 in regulating lesion innervation and pain.

Our results are consistent with prior studies that reported correlations between lesion innervation and patient pain^39–41^. Our study replicates this finding and extends it by identifying a specific subpopulation of small-diameter GFRα1-positive nociceptors (human; **Figs. 6**) and corresponding IB4-positive nociceptors (mouse; **Fig. 4**) that innervates lesions (**Figs. 2, 5, 6**) suggesting selective recruitment of nociceptors that express Ret. While this identifies a neuronal type in lesions, the functional properties of these neurons have previously not been explored.

We demonstrate in the mouse model that lesion-innervating DRG neurons are hyperexcitable compared to control neurons (**Fig. 2**). There are two notable explanations for the hyperexcitability of lesion-innervating DRG neurons. The first is that the inherent properties of nociceptors that innervate lesions render these neurons more excitable compared with control neurons. Since IB4 binding in DRG neurons specifically marks the subpopulation of Ret-positive nociceptors, we used a Ret-reporter mouse to compare the excitability of Ret+ neurons to Ret-neurons to determine if there are inherent differences in excitability between these populations^74^. We found that Ret+ neurons have a higher rheobase (less excitable) compared to Ret-neurons (**Supp. Fig. 3**), consistent with a prior report which investigated the excitability of IB4+ vs. IB4-small-diameter neurons^91^. In contrast to these findings, lesion-innervating neurons that are frequently IB4+ (**Fig. 4**), have a significantly lower rheobase (more excitable) compared to Wall-innervating neurons. These data therefore favor a second interpretation, that lesion-innervating neurons become sensitized by the highly inflamed microenvironment of lesions after recruitment. This sensitization might be through GDNF directly^58^ and/or other pathways such as tumor necrosis factor (TNF), interleukin-6 (IL-6) and interferon (IFN) signaling which are enriched in lesions from patients who report higher pain (**Supp. Fig. 4**). Although inflammatory/sensitizing factors would increase the excitability of these neurons, they would not change their identity (i.e. small-diameter nociceptors), an effect that may be evident in AP waveform similarities between Ret+ and Lesion-innervating neurons (**Supp. Fig. 3** and **Fig. 2**, respectively). We therefore propose a model in which GDNF in lesions dictates the recruitment of GFRα1-positive nociceptors, which are subsequently sensitized by inflammatory mediators known to be present in lesions, driving clinical pain.

Together, the identification of GDNF and GFRα1 positive axons in human lesions suggests that therapeutics that either 1) target excitability of sensory neurons^46^, and specifically the Ret+ subpopulation of nociceptors, or 2) block the innervation of lesions by Ret+ nociceptors, may represent viable strategies for addressing endometriosis-associated pain. Indeed, targeting GDNF signaling offers a distinct advantage as a disease modifying therapy particularly to reduce post-surgical recurrence of endometriosis pain by potentially preventing hyper-innervation of lesions by this nociceptor population. Although functional studies are required to validate this approach, clinical trials utilizing similar growth factor ligand blockade for other diseases strengthen the feasibility of the strategy^92^.

There are several limitations to our study. Animal models of endometriosis are useful tools but this disease cannot be exactly recapitulated in this model organism because the tested mice (C57Bl/6) do not menstruate, and they have a reproductive tract which does not allow for the prevailing hypothesis of the source of endometrial-like cells in the peritoneum in endometriosis^7,93^. It is interesting that we found more pain-like behaviors in Endo animals in metestrus compared to diestrus because similar cycle-dependent pain is observed in many patients (**Supp. Fig. 1**)^14^. Further, we found increased levels of components of sex-hormone signaling pathways in lesions from patients with high pain scores in our human proteomic dataset (**Supp. Fig. 4**). Investigations which explore cycle-related differences may better support the relevance of the mouse model to the human disease and encourage mechanistic testing of sex-hormone levels in relation to pain-like behaviors. While lesions that form in the mouse are consistent with some types of lesions seen in human endometriosis (**Fig. 1**), they do not represent all histologic forms of the disease^62,63^. Given the relatively early timeframe of animal experiments compared to human disease, which is often not diagnosed for years after symptom onset^7^, the mouse model lesions are most concordant with superficial peritoneal endometriosis. This aligns with human endometriosis disease trajectory where superficial peritoneal endometriosis is the dominant histology in adolescents (i.e. early in the disease)^94,95^. Regardless, GDNF expression in the gland-stromal compartment and innervation of endometrial glands in both species were similar (**Fig. 5, Supp. Fig. 5**). Another limitation is that the scope of this study is primarily restricted to lesions and lesion-innervating neurons, despite data indicating that other anatomical sites (e.g. eutopic endometrium, peritoneal fluid) are also affected by the disease^26,30,76,96–100^. While we restricted our analyses to lesions and innervating neurons, a strength of this study is that we observed that peritoneal wall innervating neurons from Endo animals exhibited a trend toward a more depolarized resting membrane potential compared to Sham littermate controls (**Fig. 2**). This finding is consistent with prior clinical data that found higher transcript levels of nociceptor-associated ion channels (TRPV1, TRPA1) in the peritoneal wall of patients with endometriosis compared to healthy controls^98^. These data suggest that neuronal sensitization may not be restricted to lesion-innervating populations but may instead reflect broader changes to the peritoneum and/or to the peripheral nervous system. Such widespread alterations could contribute to the diffuse and persistent nature of pelvic pain in endometriosis^43,44,101^ and highlight the importance of considering both lesion-specific and system-level mechanisms. Finally, the clinical sample size evaluated in this dataset (N=12 proteomics, N=6 IHC) is limited to pilot study observations. Endometriosis is a highly heterogenous disease and a larger dataset is necessary to confirm the correlation between GDNF expression level and clinical pain, and to control for factors which might alter GDNF levels such as medications, menstrual cycle and age^16^.

The results of this study are first-in-kind linking functional data from an animal model of the disease with neuronal identification in clinical, pain-defined samples. The data indicate that the Ret-positive subpopulation of nociceptors might be actively recruited into lesions through a stromal cell GDNF and sensory neuron GFRα1 signaling axis. We demonstrate that the neurons innervating endometriosis lesions in mice are highly excitable and are a distinct neuronal population of nociceptors. In addition, higher levels of GDNF and GFRα1 are associated with higher levels of patient-reported pain, connecting the observed molecular features of the lesion microenvironment to neuronal function and clinical symptoms. Together, these findings define a mechanistic framework for how lesion innervating DRG neurons contribute to endometriosis-associated pain and identify lesion GDNF and GFRα1-positive nociceptors as tractable targets for therapeutic intervention.

## Methods

### Animal model

Adult female C57Bl/6J mice (Jackson Laboratory, Stock #000664) were utilized in all studies due to the nature of the disease impacting persons with a uterus. Animals were housed on a 12-hour light/dark cycle with free access to food and water. To induce the endometriosis model, a syngeneic donor-host model was utilized as previously described^47^. Briefly, five-week-old donor mice were primed with a subcutaneous injection of 3 µg estradiol benzoate (Cayman Chemical, #10006487) dissolved in 100% ethanol and brought to 100 µL in sterile saline seven days prior to tissue harvest. Following euthanasia, the uterine horns of the donor mice were harvested and longitudinally incised to expose the endometrium. The tissue was minced into fragments (<1 mm^3^) in ice-cold Hanks’ Balanced Salt Solution (HBSS). The resulting fragments were subsequently resuspended in 500 µL of pre-warmed (37°C) HBSS. Littermate recipient animals were randomly assigned to either the endometriosis (Endo) or Sham group. Endo mice received an intraperitoneal (i.p.) injection of the uterine tissue suspension (one donor horn per recipient) via a 1 mL syringe fitted with an 18-gauge needle. Sham animals received 500 µL of warm HBSS without tissue. To mitigate cage-effect bias and ensure investigator blinding in subsequent assays, each cage housed both Sham and Endo animals as littermate controls.

### Behavior

All behavioral testing was performed by an investigator blinded to the experimental groups. At five and eight weeks post-disease induction, animals were acclimated to a raised wire mesh flooring within individual transparent plexiglass chambers for 30 minutes prior to assessment. Ambient room conditions were kept consistent throughout all studies, and all testing was conducted in the morning. First, animals were observed without disruption for 10 minutes to quantify spontaneous pain-like behaviors. A behavioral bout was defined as the continuous episode of one of the following specific behaviors from onset to cessation: “Abdominal squashing” was defined as the animal pressing its abdomen against the mesh floor as previously described^47^; Writhing was defined as abnormal stretching or rotation of the abdomen, consistent with models of visceral pain^102^; Abdominal licking was defined as grooming behaviors specifically directed to the abdomen that was not subsequent to other grooming behaviors. Following the assessment of spontaneous behaviors, abdominal mechanical sensitivity was measured by percentage withdrawal response to von Frey filaments of increasing force (0.04, 0.08, 0.16, and 0.32 grams). This testing paradigm was chosen based on pilot data showing a dynamic range of responses in control animals across the forces, with ceiling or floor effects observed outside this range (data not shown). Positive responses were defined as the animal retreating from the filament, kicking the hind legs, or jumping in response to the fiber. The number of positive responses out of 10 total trials per filament was recorded to calculate the percent response. An inter-stimulus interval of 1-2 minutes was maintained between applications, with at least five minutes between different filaments. At the end of behavioral assessments, the external genitalia were observed and each animal was assigned to an estrous cycle phase (proestrus, estrus, metestrus, or diestrus) based on previous reports^66^. Finally, animals were weighed and returned to their home cages.

### Tissue Collection and Histological Validation

For gross anatomical characterization of the disease, animals were euthanized by cervical dislocation under deep isoflurane anesthesia. The peritoneum was carefully opened, and the abdominal walls were systematically surveyed for the presence of endometriotic-like lesions. Next, the reproductive organs, abdominal fat pads, and mesentery were inspected for additional ectopic lesions. The location and number of lesions were documented and biopsies were embedded and snap-frozen in Optimal Cutting Temperature (OCT) compound on dry ice. To confirm the presence of endometrial glands and stroma, histological validation was performed on 20 µm cryosections. Briefly, sections were stained with Mayer’s Hematoxylin (Sigma-Aldrich) and Eosin (Sigma-Aldrich) (H&E) according to standard protocols, followed by ethanol dehydration.

### Retrograde dye injections

At eight weeks post-disease induction animals underwent *in vivo* retrograde labeling of DRG neurons that innervate endometriotic lesions or the peritoneal wall. Under isoflurane anesthesia (2.5% and 2% oxygen), a midline laparotomy was performed to expose the peritoneal cavity and visualize the abdominal wall. Fluorophore-conjugated Wheat Germ Agglutinin (WGA; WGA-640R #29026-1 and WGA-532 #29064-1; Biotium) were reconstituted in sterile ultrapure water and stored at -20°C. For targeted injections, either an endometriotic lesion or control peritoneal wall tissue was pierced with a 33-gauge needle attached to a Hamiliton syringe and allowed to equilibrate for 1 minute. 0.5-2 µL of WGA was injected into the tissue over 60 seconds, with volume dependent on the capacity of the tissue. After injection, the needle remained in the tissue for at least one minute prior to slow retrieval to minimize tracer backflow. As the needle was withdrawn, sterile gauze was immediately placed on the site to prevent leak of the dye. In a subset of animals, distinct WGA fluorophores were injected into different anatomical locations (e.g. lesion vs. peritoneal wall) to distinguish specific innervation patterns. Following injections, the abdominal muscles were sutured with 6-0 silk, and the skin was approximated with surgical staples. Animals were allowed to recover for 3-7 days to permit optimal retrograde transport prior to tissue harvest.

### DRG Dissociation and Electrophysiology

Following retrograde tracer transport, animals were euthanized and transcardially perfused with ice-cold HBSS. Bilateral dorsal root ganglia (DRG) from spinal levels T10 through S2 were rapidly dissected and placed into ice-cold HBSS. Immediately following DRG collection, gross disease pathology was confirmed as described above. DRG neurons were dissociated as previously described^68^. Briefly, ganglia underwent enzymatic digestion with papain (0.33 mg/mL; Worthington) for 20 minutes at 37°C, followed by collagenase type II (1.5 mg/mL; Sigma-Aldrich) for 20 minutes at 37°C. After enzymatic treatment, cells were mechanically triturated, washed, and passed through a 40-µm cell strainer. Cells were centrifuged at 1000 rpm for 3 minutes, resuspended, and plated onto glass coverslips pre-coated with poly-D-lysine and collagen. Neurons were maintained in complete DRG medium consisting of Neurobasal A (Gibco) supplemented with 5% fetal bovine serum (Gibco), 1% penicillin/streptomycin (Corning), GlutaMAX (Life Technologies), and B-27 supplement (Gibco).

Whole-cell patch-clamp recordings were performed 16-48 hours post-plating, with recording times matched across groups. Recording parameters were implemented as previously described^68^ at room temperature. Recordings were performed in an external solution containing 145 mM NaCl, 2 mM CaCl2, 1.2 mM MgCl_2_, 7 mM glucose, and 10 mM HEPES, pH 7.3 with NaOH and 300-310 mOsm. Cells were recorded within 1 hour of removal from DRG media. Neurons were required to have a stable resting membrane potential (RMP) <-35 mV and stable access resistance. To visualize WGA fluorophores, a 625 or 530 nm LED light sources (ThorLabs) were used. Once a neuron was identified to be labeled by a single dye, a thick-walled borosilicate glass recording pipette (Sutter Instrument) with an average resistance of 4-6 MΩ (pulled with a P-97 horizontal puller; Sutter Instrument) containing intracellular solution (120 mM potassium gluconate, 5 mM NaCl, 2 mM MgCl_2_, 0.1 mM CaCl_2_, 10 mM HEPES, 1.1 mM EGTA, 4 mM Na_2_ATP, 0.4 mM Na_2_GTP, 15 mM sodium phosphocreatine, adjusted to pH = 7.3 with KOH, and 292 mOsm with sucrose) was used to create a giga-ohm seal. Data were acquired using a MultiClamp 700B amplifier and a Digidata 1550B digitizer (Axon Instruments) controlled by Clampex software (v11.1; Molecular Devices). Signals were sampled at 20 kHz and analyzed offline.

Upon achieving the whole-cell configuration, intrinsic and evoked electrophysiological properties were recorded in current-clamp mode. Intrinsic properties included cell diameter, membrane capacitance, spontaneous activity and RMP. To assess evoked excitability, cells were held at - 60 mV and stimulated with a series of 1 second depolarizing step stimuli (square pulses) at 10 pA increments. Rheobase was defined as the minimum amount of current necessary to evoke at least one action potential (AP). To evaluate repetitive firing capacity, current was injected at multiples (1-4x) of the calculated rheobase. Neurons were classified as repetitive firing if they fired more than one AP during any of these current injections. AP kinetics were analyzed from the first AP following each cell’s rheobase. These parameters include the AP^68^: threshold (voltage when the first derivative of the potential exceeded 20 mV/ms), half-width (time at 50% AP amplitude), amplitude (voltage difference from threshold to peak), and AP peak (maximum depolarized membrane potential reached during the AP). All electrophysiological data were analyzed offline using Easy Electrophysiology software (v2.6.1).

### Immunohistochemistry

All tissue was collected fresh and snap frozen on dry ice. DRG were sectioned at 10 µm and non-neuronal tissue was sectioned at 20 µm using a cryostat. Sections were mounted onto slides to be used immediately or stored -20°C until use^36^. A perimeter around sections was drawn using a hydrophobic pen (Vector Laboratories) prior to fixing with 4% paraformaldehyde (PFA) for 10 minutes at room temperature. After washing with PBS, tissue was blocked and permeabilized with a buffer containing 1% BSA in PBS, 0.1% Triton-X 100 and 0.1% Sodium Azide for one hour at room temperature before incubating with the target antibody/s overnight at 4°C in blocking buffer. Primary antibodies included: Chicken anti-Peripherin (1:500 or 1:1000; Part#: A21449); goat anti-GDNF (1:100; R&D Systems; Part#: AF-212-NA); and rabbit anti-GFRα1 (1:100; Abcam; Part#: Ab8026). The next day, the slides were washed in PBS prior to appropriate fluorescent-conjugated secondary antibody incubation (1:500) and/or Isolectin B4 (1:300; Invitrogen; Part#:121412; in the presence of 3mM CaCl_2_ and MgCl_2_) for one hour at room temperature. Slides were washed prior to counterstaining with DAPI (1:10,000; Invitrogen; #D1306) and coverslipping with Prolong Gold Antifade Mountant (Invitrogen). Images were captured using a confocal microscope (Leica Stellaris 5). Acquisition parameters were kept consistent across groups by an investigator blind to the groups. All histological analyses were completed by an experimenter blind to the condition and pain scores. For lesion analysis, regions of interest (ROIs) were manually drawn around all glands in tissue sections identified by characteristic epithelial and stromal layers. The mean fluorescence intensity (MFI) was then calculated for each ROI across at least three nonconsecutive sections per sample, given that glands were identified. To account for variations in stromal abundance within lesions, the average GDNF MFI was normalized to the average DAPI signal per participant as an approximation of cell density. Correlative analysis between Peripherin and GFRα1 was performed by relating the average MFI per participant. To assess the overall pathway activity, the MFI for Peripherin, GFRα1 and GDNF were converted to standardized Z-scores by the following equation: Z_Factor_ = (MFI_Factor_ – MFI_Factor Mean_) / MFI_Factor Standard Deviation_. The Z-score for each factor was summed to create a single score for each participant (Composite Z-Score)^83,103^.

### Proteomics

Total protein was isolated from surgically identified lesions (5-30 mg). Protein was isolated in 350 µL RIPA buffer (Sigma-Aldrich, #R0278) containing a protease inhibitor cocktail (Sigma-Aldrich, #11836153001) using a motorized probed micro-tissue homogenizer on ice prior to centrifugation at 14,000 *g* for 10 minutes at 4°C. The supernatant total protein was calculated using a BCA assay according to manufacturer’s direction (Thermo Scientific, #23225). Samples were diluted to 0.5 mg/mL total protein and analyzed using the OLink Proteomic Services (Explore HT) technologies at the High-Throughput Biomarker Core at Vanderbilt University Medical Center using next generation sequencing (NGS) and including quality control, NGS read counts, and Normalized Protein Expression (NPX) values. NPX is a relative quantification unit on a log_2_ scale. This data was exported into R (v4.3.3) for downstream analysis with the OlinkAnalyze (v4.3.1), limma (v3.56.2), and ggplot2 (v3.5.2) packages. Quality control was assessed through principal component analysis (PCA), and differential expression between high/severe and mild/moderate groups was tested with the empirical-Bayes moderated t-statistic from limma. Growth factors were defined *a priori* as a curated list of 52 genes covering the GDNF/neurotrophin family (GDNF, BDNF, NGF, NTF3/4, CNTF, NRTN, ARTN, PSPN), VEGF, PDGF, FGF, EGF, IGF/IGFBP, TGF-β, BMP, NOTCH1 and colony-stimulating-factor families. Per-protein effect sizes were summarized as forest plots with family-wide Benjamini-Hochberg correction. Pathway enrichment was performed on the differential-expression (DE) result from the limma high/severe vs. mild/moderate contrast, using the R package clusterProfiler with org.Hs.eg.db for human gene annotation, ReactomePA for Reactome pathway enrichment, and enrichplot for visualization.

### Patient data

Participants were recruited through the Obstetrics and Gynecology (OBGYN) clinics at Washington University and Barnes-Jewish Hospital (**Table 1**). Prior to scheduled minimally invasive gynecological surgery for the diagnosis and resection of endometriosis lesions, participants completed a series of surveys during a pre-surgical study visit. The clinical pain metrics were used for all subsequent correlations with proteomics and histological data. The pain surveys included ratings on the average intensity of endometriosis-related pain over the preceding 30 days on a scale ranging from no pain to the worst pain imaginable. In addition, participants completed the Endometriosis Health Profile 30 (EHP-30) questionnaire^104^ which assesses self-reported quality of life of women with endometriosis within the past 4 weeks. The survey includes 30 multiple choice questions, and the sum of the questions was calculated with higher scores indicating a worse quality of life. In the present study, the pain subscale was used with higher scores indicating worse pain^105^. After the surgery, patients were clinically assigned endometriosis based on the standards of the American Association of Gynecologic Laparoscopists (AAGL) scoring system which evaluates surgical complexity based on the number, location and infiltration depth of lesions^31^.

## Statistics

Statistical analyses and data visualization were performed using R v4.3.3 or GraphPad Prism v10.3.1. Normality of the data was assessed using the Shapiro-Wilk test. Measurements of two groups over time from the same animal were analyzed using a two-way repeated measures ANOVA followed by Tukey’s post hoc analysis. For data involving two groups over time from different animals, a two-way ANOVA with Tukey’s post hoc analysis was utilized. Measurements between three or more groups at a single timepoint were tested via one-way ANOVA followed by Tukey’s post hoc test, or the Kruskal-Wallis test with Dunn’s post hoc analysis for non-normally distributed data. Categorical data were analyzed using Fisher’s exact test. Linear regressions were performed to measure associations between multiple proteins or between a single protein and clinical pain scores. The goodness of fit (R^2^) and the result of the F-test result are presented for each association. Appropriate tests of variation are denoted in individual figure legends. The critical significance value was set at α<0.05, and exact p values and associations are marked on figure panels or in individual figure legends.

## Study Approval

### Animals

All experimental procedures were approved by the Institutional Animal Care and Use Committee (IACUC) of Washington University in St. Louis and conducted in accordance with the US National Institutes of Health (NIH) Guide for the Care and Use of Laboratory Animals.

### Patients

All study procedures were approved by the Washington University Institutional Review Board and written informed consent or assent was obtained from all participants prior to enrollment. The study was preregistered in ClinicalTrail.gov (NCT06101303). Inclusion criteria included patients aged 12-45 years old with suspected or known endometriosis. Exclusion criteria included pregnancy and lactation.

## Data Availability

Data is available in the Supporting Data Values file.

## Author Contributions

A.J.D. conceived of and performed most of the experiments and analyses and wrote the manuscript. M.F. helped to establish the mouse model including behavioral and confirmation/cycle related analyses including histology. A.J.K. helped perform and analyze human immunohistochemistry levels. J.M.M. performed proteomic analyses. M.E.M. performed Ret/IB4 *in vitro* experiments. R.B. helped perform mouse immunohistochemistry analyses.

J.G.P. advised and helped with the Ret/IB4 experiments. E.B. and W.T.R. are gynecologic surgeons who performed the endometriosis resection surgeries. W.T.R. and H.N.A. oversaw recruitment of patients to the study. W.T.R, H.N.A. and R.W.G. helped conceive of the project and made primary edits to the manuscript.

## Funding Support

National Institute of General Medical Sciences (NIGMS), Washington University School of Medicine, Department of Anesthesiology training grant T32GM108539 (AJD). National Institute of Health, National Institute of Child Health and Human Development, 1R21HD115568-01 (HNA/WTR). National Institute of Health, National Institute of Child Health and Human Development, 1K23HD110710-01 (WTR).

## Notes

### Competing Interest Statement

The authors have declared no competing interest.

