## Supplemental Figures for "Preferential innervation of endometriosis by hyperexcitable Ret/GFRα1+ nociceptors associates with target GDNF and clinical pain"

**Supplementary Materials**

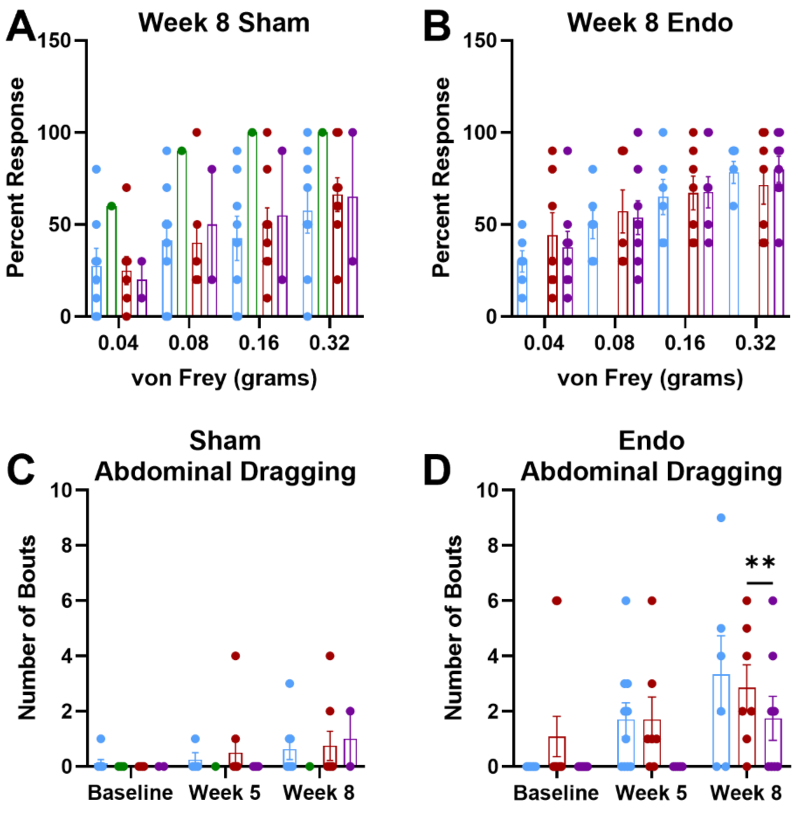


**Supplemental Figure 1. The effect of estrous cycle on Endo model pain-like behaviors.** Animals were classified into four stages of the estrous cycle by visual inspection of external genitalia. At eight weeks we found no difference in mechanical sensitivity between estrous stage in Sham animals **(A)** or in Endo animals **(B)**. **C.** Similarly, there was no difference in the number of bouts of abdominal dragging in Sham animals. **D.** However, there was a statistically significant difference in the number of abdominal dragging bouts in Endo animals between the metestrus and diestrus phase (**p=0.0074). Two way ANOVA, Tukey's; Sham n= proestrus 8/4/8, estrus 3/1/1, metestrus 8/10/8, 2/4/2, baseline/week 5/week 8, respectively. Endo n= proestrus 6/10/6, estrus 0/0/0, metestrus 11/7/7, diestrus 7/5/8, baseline/week 5/week 8, respectively. Data represented as mean +/- SEM.

A

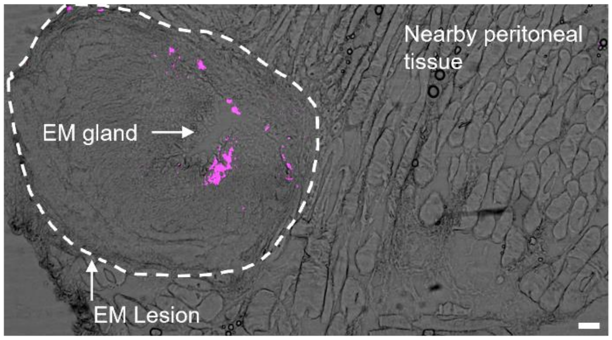


B

| | Amplitude (mV) | Threshold (mV) | Half width (ms) | Input resistance (MOhms) | Capacitance (pF) |
|---|---|---|---|---|---|
| Sham Wall (n=11) | 64.48 +/- 3.3 | -9.39 +/-2.3 | 2.87 +/- 0.26 | 271.4 +/- 44.98 | 38.8 +/- 4.66 |
| Endo Wall (n=11-13) | 60.50 +/- 4.5 | -2.34 +/- 6.3 | 3.00 +/- 0.45 | 250.5 +/- 48.98 | 52.34 +/- 8.32* |
| Lesion (n=17) | 51.48 +/- 5.1 | -11.42 +/- 3.5 | 3.12 +/- 0.34 | 335.5 +/- 48.48 | 31.38 +/- 3.49 |

**Supplemental Figure 2. Tissue dye visualization and additional electrophysiological readouts. A.** Representative peritoneal wall section containing an endometriosis lesion in mouse. Purple staining demonstrates retrograde dye localization to the lesion without spread to the nearby muscle layers of the abdominal wall. Annotations describe features of the lesion and tissue. Scale=50 µm. **B.** Electrophysiological data indicating no differences between lesion-innervating groups, wall innervating groups from animals with Endo, or wall innervating groups from Sham animals for AP amplitude, threshold, half width or input resistance. There was a significant increase in the capacitance of Endo Wall neurons relative to lesion-innervating neurons (One way ANOVA, Tukey’s or Kruskal-Wallis, Dunn’s as appropriate; *p=0.021 vs. Lesion; n=11-17/group)**.** Data represented as mean +/- SEM.

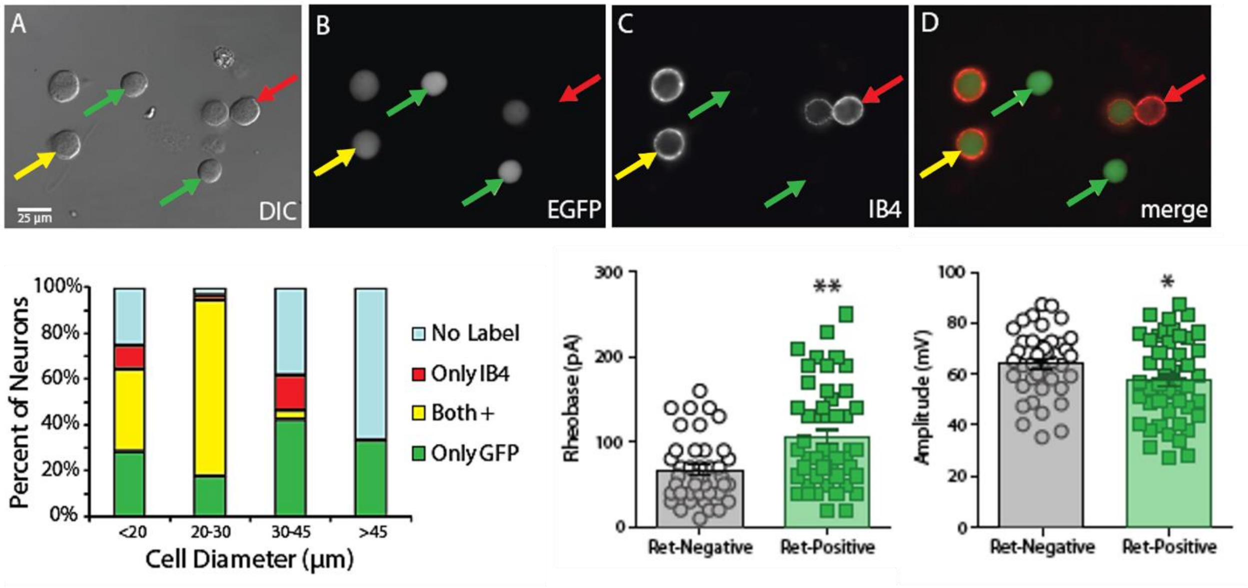


| | Peak (mV) | Threshold (mV) | Half width (ms) |
|---|---|---|---|
| Ret-Negative | 45.58 | -18.54 | 1.72 |
| Ret-Positive | 40.67$^{\#}$ | -16.84 | 1.88 |

**Supplemental Figure 3. Small-diameter Ret+ DRG neurons are IB4+ with distinct electrophysiological properties. A-D.** Representative images of DRG neurons *in vitro* from animals which genetically express EGFP (Ret$^{EGFP/+}$) and stain positive or negative to IB4 (yellow arrow = EGFP+ and IB4+; green arrow = EGFP+ alone; red = IB4+ alone). **E.** Proportional analysis of all possible groups by neuron diameter revealing that small-diameter neurons (≤30 µm) are primarily both IB4+ and Ret+. **F.** Patch clamp electrophysiology of small-diameter neurons reveal that Ret+ neurons have a higher rheobase and (**G**) have a lower AP amplitude. **H.** Additional electrophysiological properties between Ret+ and Ret- small-diameter neurons reveal a trend toward a lower AP peak in Ret+ cells (#p=0.065 vs. Ret negative cells), with no differences in threshold or half width. Student t test; *p<0.05, **p<0.01.

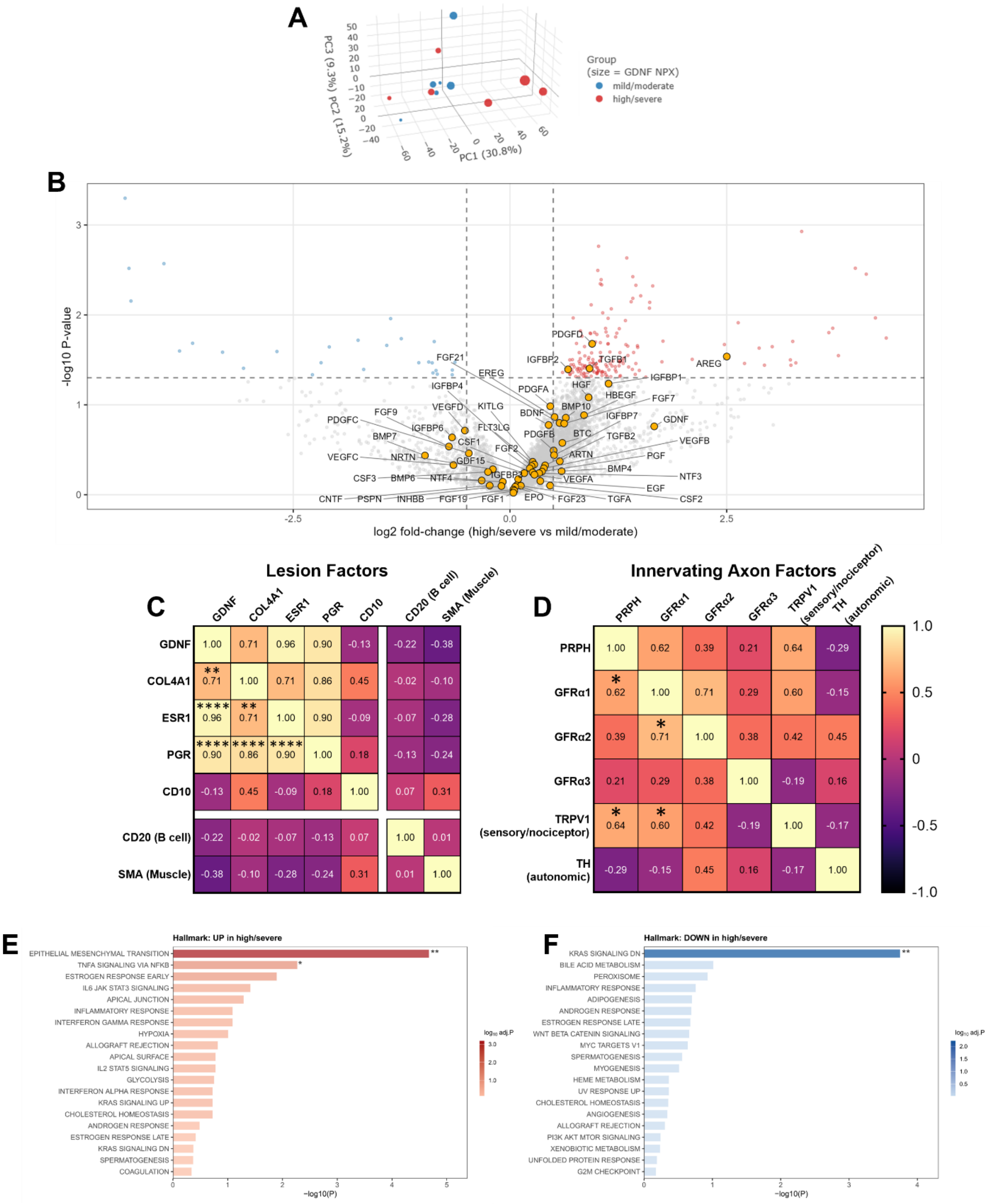


**Supplemental Figure 4. Proteomic enrichment analysis and between factor associations.**

**A.** Principal component analysis (PCA) of samples divided by pain intensity (red = high/severe pain, blue = mild/moderate pain) as well as relative GDNF expression (size of dots). **B.** Volcano plot indicating factors which are non-significantly and significantly enriched or dis-enriched in lesions from high/severe pain reporting patients (Intensity = 7-9; N=6) compared to patients who reported mild/moderate pain (Intensity = 3-6; N=6). Select growth factors are marked. **C.** Proteomic data associations indicating correlations between GDNF and various markers of stromal cell or endometriosis stroma, but not with other cell types including B cells and muscle cells. **D.** Similar correlation matrix indicating associations between Peripherin and DRG neuron subtypes by growth factor receptor expression indicating an association between Peripherin and GFRα1 but not GFRα2 and GFRα3. TRPV1 levels, which is restricted to sensory neuron nociceptors, also correlate with GFRα1 levels. Peripherin and TRPV1 are related, but Peripherin and TH, which is restricted to autonomic fibers, are not. Pearsons r is indicated on correlation matrix heatmaps, asterisks indicate statistical significance. **E-F.** Pathway enrichment analysis indicates up and down regulated signaling pathways between pain-groups (*p<0.05, **p<0.01, ****p<0.0001).

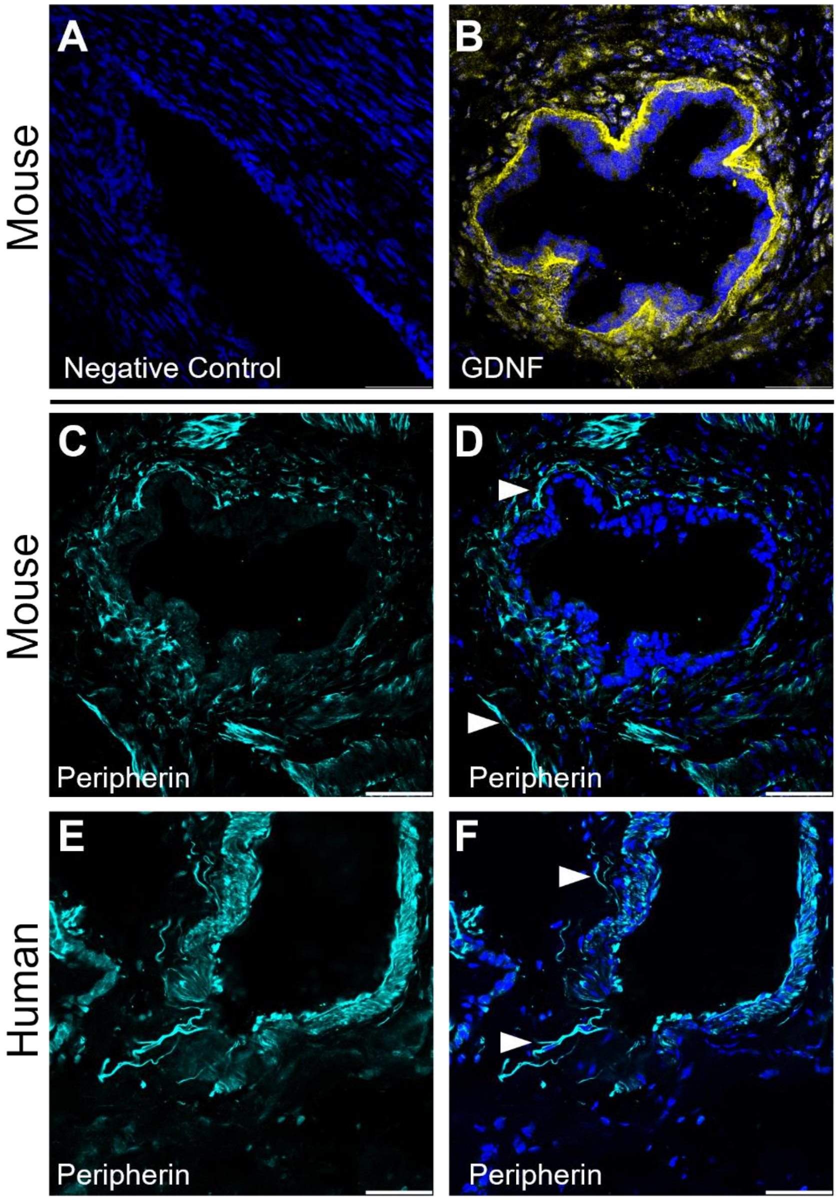


**Supplemental Figure 5. Comparisons of lesion GDNF in mouse as well as innervation patterns between mouse and human endometrial-like glands. A-B.** Images indicating GDNF staining in mouse lesions similar to human lesions. **C-D.** Representative images of mouse peripherin in endometriosis and (**E-F**) human peripherin in endometriosis. Arrows indicate axons entering and within the stromal layer of endometrial glands. Scale=50 µm.

| **Additional IHC correlations with VAS** | **GDNF dataset** | **GFRα1/Peripherin dataset** |
|---|---|---|
| Average DAPI | $R^2$=0.296, p=0.26 | $R^2$=0.203, p=0.37 |
| Average Area | $R^2$=0.018, p=0.80 | $R^2$=0.007, p=0.88 |

**Supplemental Table 1. Cell density and area measured are not associated with patient reported pain (VAS).** Goodness of fit and significance values indicated for DAPI (an approximation of cell density) in the measured portion of endometrial glands in both IHC datasets, as well as the total area measured around endometrial glands for each dataset. No associations were found.
